# Impact of maternal malnutrition on gut barrier defence: implications for pregnancy health and fetal development

**DOI:** 10.1101/635052

**Authors:** Sebastian A. Srugo, Enrrico Bloise, Tina Tu- NgocThu Nguyen, Kristin L. Connor

## Abstract

Small intestinal Paneth cells, enteric glial cells (EGC), and goblet cells maintain gut mucosal integrity, homeostasis, and influence host physiology locally and through the gut-brain axis. Little is known about their roles during pregnancy, or how maternal malnutrition impacts these cells and their development. Pregnant mice were fed a control diet (CON), undernourished by 30% vs. control (UN), or fed a high-fat diet (HF). At day 18.5 (term=19), gut integrity and function were assessed by immunohistochemistry and qPCR. UN mothers displayed reduced mRNA expression of Paneth cell antimicrobial peptides (AMP; *Lyz2, Reg3g*) and an accumulation of villi goblet cells, while HF had reduced *Reg3g* and mucin (*Muc2*) mRNA and increased lysozyme protein. UN fetuses had increased mRNA expression of gut transcription factor *Sox9*, associated with reduced expression of maturation markers (*Cdx2, Muc2*), and increased expression of tight junctions (TJ; *Cldn-7*). HF fetuses had increased mRNA expression of EGC markers (*S100b, Bfabp, Plp1*), AMP (*Lyz1, Defa1, Reg3g*), and TJ (*Cldn-3, Cldn-7*), and reduced expression of an AMP-activator (*Tlr4*). Maternal malnutrition altered expression of genes that maintain maternal gut homeostasis, and altered fetal gut permeability, function, and development. This may have long-term implications for host-microbe interactions, immunity, and offspring gut-brain axis function.

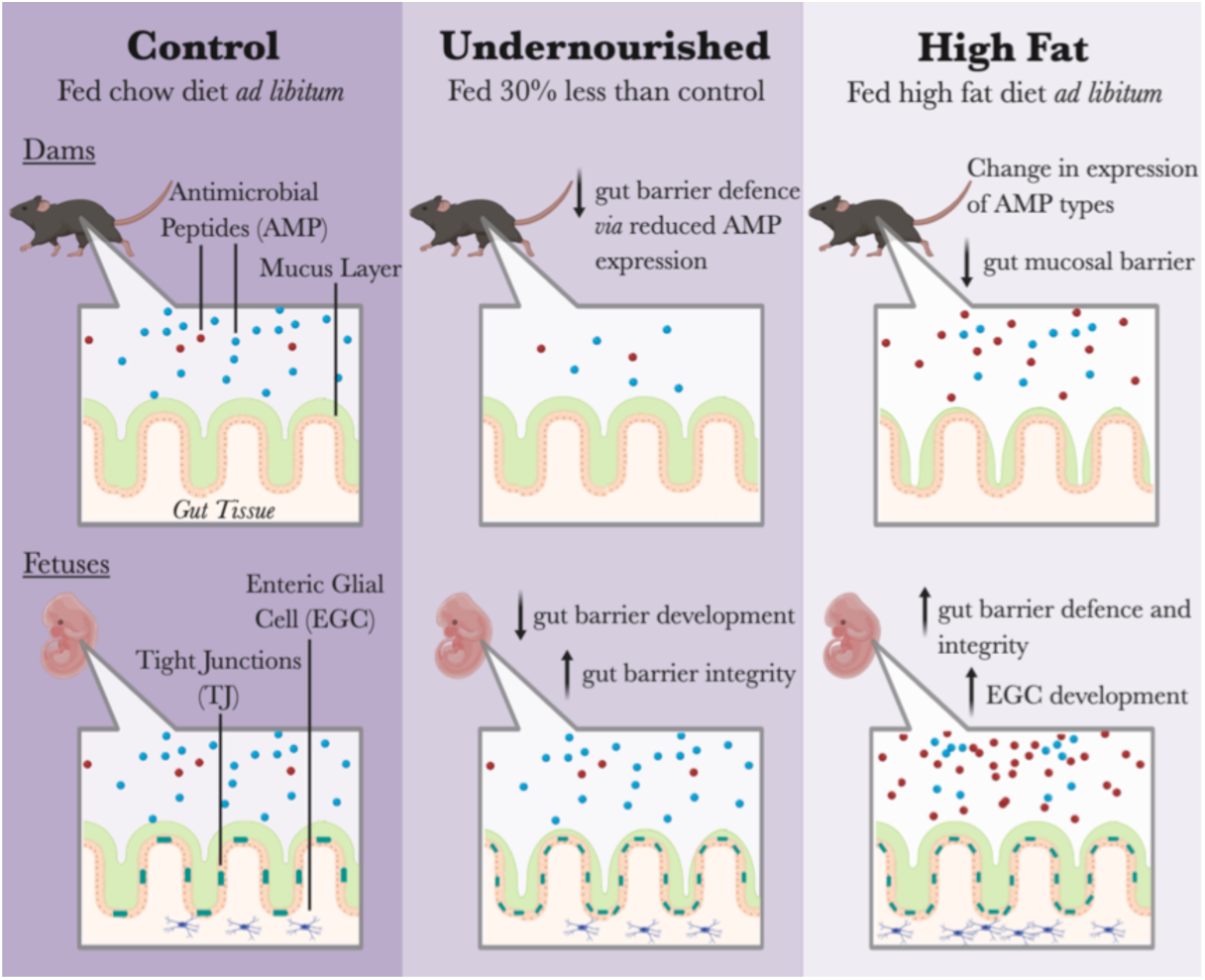

## Introduction

The gut is critical to host health and disease development through interactions with microbes that colonize the gut and establishment of the gut epithelial barrier (1,2). In health, the gut is involved in vitamin synthesis (3), nutrient metabolism (4), and protection against pathogens (1,5). It is also important for brain function: a direct and bidirectional channel of communication exists between the gut and the brain, dubbed the gut-brain axis, which connects the enteric and central nervous systems (CNS) (6). Microbes that reside in the gut, their metabolites (7), hormones (8), and immune factors (9), have been shown to affect gut and brain function through this axis in humans and animals (6). Since microbes and the metabolites they produce can activate host immune, metabolic, and stress pathways once they leave the gut environment (10), an important part of the gut-microbe-host relationship is the separation between the gut tissue, the gut ecosystem, and the rest of the host.

This separation is mediated by the intestinal epithelial barrier, which lies at the interface between exogenous host factors and the internal gut microenvironment and helps to regulate microbe-host interactions. During periods of optimal nutrition and host health, two key cell types are involved in supporting and maintaining this gut barrier: Paneth cells and enteric glial cells (EGCs). Paneth cells reside in the epithelium of the small intestine (SI), maintain gut integrity through production of tight junction (TJ) proteins (11), and produce antimicrobial peptides (AMPs) which regulate the host-microbe relationship (12). As well, due to their proximity to crypt stem cells (13,14), Paneth cells can affect gut epithelial cell differentiation and gut maturation (13,14). Toll-like receptors (TLRs), which activate the innate immune response when bacterial components are recognized in the gut environment, are purported to play a key role in inducing Paneth AMP production (15). In addition, EGCs are part of the enteric nervous system (ENS) and gut-brain axis (16), and respond to and control gut inflammation (17,18). EGCs also influence gut integrity and permeability (19) through their long cytoplasmic processes which make direct contact with the barrier (20). Other enteric cells, such as goblet cells, maintain gut barrier integrity through their production of mucus (11). Importantly, Paneth cells and EGCs are established in early development (Figure S1) (13,21), suggesting they are key for early and lifelong gut and brain health.

The integrity of the gut barrier and the composition and function of the gut microbiome can be greatly affected by the diet of the host (30–32), since the indigestible polysaccharides in the host diet become gut bacterial substrates and nutrients (33). Indeed, adaptations to gut bacterial metabolism and transcription occur within days of dietary changes in humans (34). In response to both over-(35) and undernutrition (36), collectively known as malnutrition, gut barrier function and integrity can become dysregulated, leading to gut microbial dysbiosis, altered gut function, and a leaky gut barrier. Yet, few studies have elucidated the effects of malnutrition on Paneth cell (37–41) and EGC (42,43) development and function (indeed, none have examined EGCs during undernutrition to our knowledge), or how either of these cells are affected by and/or perpetuate gut dysfunction. Since the rates of both undernutrition and underweight (44), as well as over-nutrition and obesity (44), are increasing worldwide, and a compromised gut barrier is both caused by, and leads to, a variety of immune-related, and chronic diseases, the effects of malnutrition on the gut-host relationship are important to understand.

During pregnancy, the intestinal epithelial barrier and the microbes contained within the gut are doubly important, as they protect both the mother and, by association, the fetus from harmful bacteria and xenobiotics (1,45), produce nutrients required for pregnancy health (46), and absorb nutrients into the blood stream that are vital to fetal development (47,48). We and others have shown that maternal malnutrition impacts the maternal gut microbiome (49–52) and is associated with increased levels inflammation in the maternal gut and peripheral circulation (49,52). In offspring, a mature intestinal epithelial barrier ensures a healthy and homeostatic gut environment (53), which allows the offspring to appropriately respond to infections (53), absorb and produce nutrients (15), and likely establish optimal communication with the brain and other organs (6). Still, little is known about how malnutrition impacts the maternal epithelial barrier during pregnancy, a ‘stress-test’ in itself, or whether maternal malnutrition adversely programmes fetal gut development and function. Additionally, although we know that Paneth cells (13) and EGCs (22,23) are laid down and functional in early life, we know less about how early life adversity, including poor nutrition, or gut microbes, may influence their development and function.

We therefore sought to answer two questions: how does malnutrition during pregnancy affect maternal gut barrier function, and does maternal malnutrition impact fetal gut integrity, function, and development? We focused on Paneth cells and EGCs to answer these questions as these cells may be critical for long term gut function and communication with the brain. We hypothesised that malnutrition would lead to an adverse maternal gut environment, and that mothers fed a high fat (HF) diet and their offspring would display the most affected gut function, since our previous work has demonstrated that the maternal gut is especially affected by a HF diet (49). We also hypothesised that, due to the changes in maternal diets and gut environments, the fetal gut would display aberrant gut integrity and function and EGC development, with different outcomes in fetuses from UN and HF mothers.

## Methods

### Animal model

All housing and breeding procedures were approved by the Animal Care Committee at Mount Sinai Hospital (Toronto, ON, Canada; AUP 16-0091H). Male and female C57BL/6J mice (Jackson Laboratories, Bar Harbor, ME, USA) were housed using a 12:12 light:dark cycle at 25°C, with free access to food and water. Females were randomised into three diet groups: mice fed control diet *ad libitum* before and throughout pregnancy (CON, n=7); mice fed control diet *ad libitum* before mating and undernourished by 30% from gestational day (d)5.5 to 17.5 (UN, n=7); and mice fed 60% high-fat diet *ad libitum* from 8 weeks prior to mating and through pregnancy (HF, n=8). Males were fed control diet *ad libitum* and mated with females at ∼10 weeks of age. Females were housed individually following confirmation of pregnancy status (presence of vaginal sperm plug). Dams were weighed weekly before and daily during pregnancy.

### Maternal SI and fetal gut collection

Dams were sacrificed by cervical dislocation at d18.5 (term=d19). Fetuses were collected, weighed, and at random, one male and female fetus from each litter was used for fetal biospecimen collections. Maternal and fetal gastrointestinal (GI) tracts were dissected as detailed previously (49). A 2– 5mm piece of maternal SI from the mid portion of the SI (representing the jejunum) and the entire fetal gut were flash frozen in liquid nitrogen then stored at −80°C for later molecular analyses. Another 2–5mm of maternal SI from the mid portion was flushed with buffered 4% paraformaldehyde (PFA), cut longitudinally, and cut into two pieces for fixing in 4% PFA at 4°C overnight. Fixed SI were washed thrice with 1x PBS and stored in 70% ethanol until paraffin embedded for later immunohistochemical analyses.

### RNA extraction and mRNA expression analysis

Total RNA was extracted from maternal SI and fetal guts using the Tissue Lyser II (Qiagen, Hilden, NRW, Germany) and RNA extraction kits following manufacturer’s instructions (QIAGEN RNeasy Plus Mini Kit, Toronto, ON, Canada). Eluted RNA quality and quantity were assessed by spectrophotometry (DeNovix, Wilmington, DE, USA), and 1ug of RNA was reverse transcribed using 5X iScript Reverse Transcription Supermix (Bio-Rad, Mississauga, ON, Canada).

We focused on genes (Table 1) involved in gut barrier function, integrity, and development to establish how maternal malnutrition may impact the maternal gut and, by consequence, fetal gut and ENS development. mRNA expression data were normalised to the geometric mean of the three stably-expressed reference genes: TATA-Box Binding Protein (*Tbp*), Tyrosine 3-Monooxygenase/Tryptophan 5-Monooxygenase Activation Protein Zeta (*Ywhaz*) and Beta-actin (*Actb*). Primers were designed from gene sequences found in the NCBI Nucleotide Database or taken from the literature and analyzed using NCBI Primer-BLAST and Oligo Calc (Northwestern University, Evanston, IL, USA) for appropriate gene targeting and properties. Amplification and detection of mRNA expression was measured using CFX384 Touch Real-Time PCR Detection System (Bio-Rad). Samples, standards, and controls were pipetted in triplicate. Inter-run calibrators and non-template controls were run alongside each gene to normalise between plates and to assess contamination, respectively. The PCR cycling conditions were: 30sec at 95°C, 40x 5sec at 95°C, 20sec at 60°C. Data were analyzed applying the Pfaffl method.

**Table 1.**
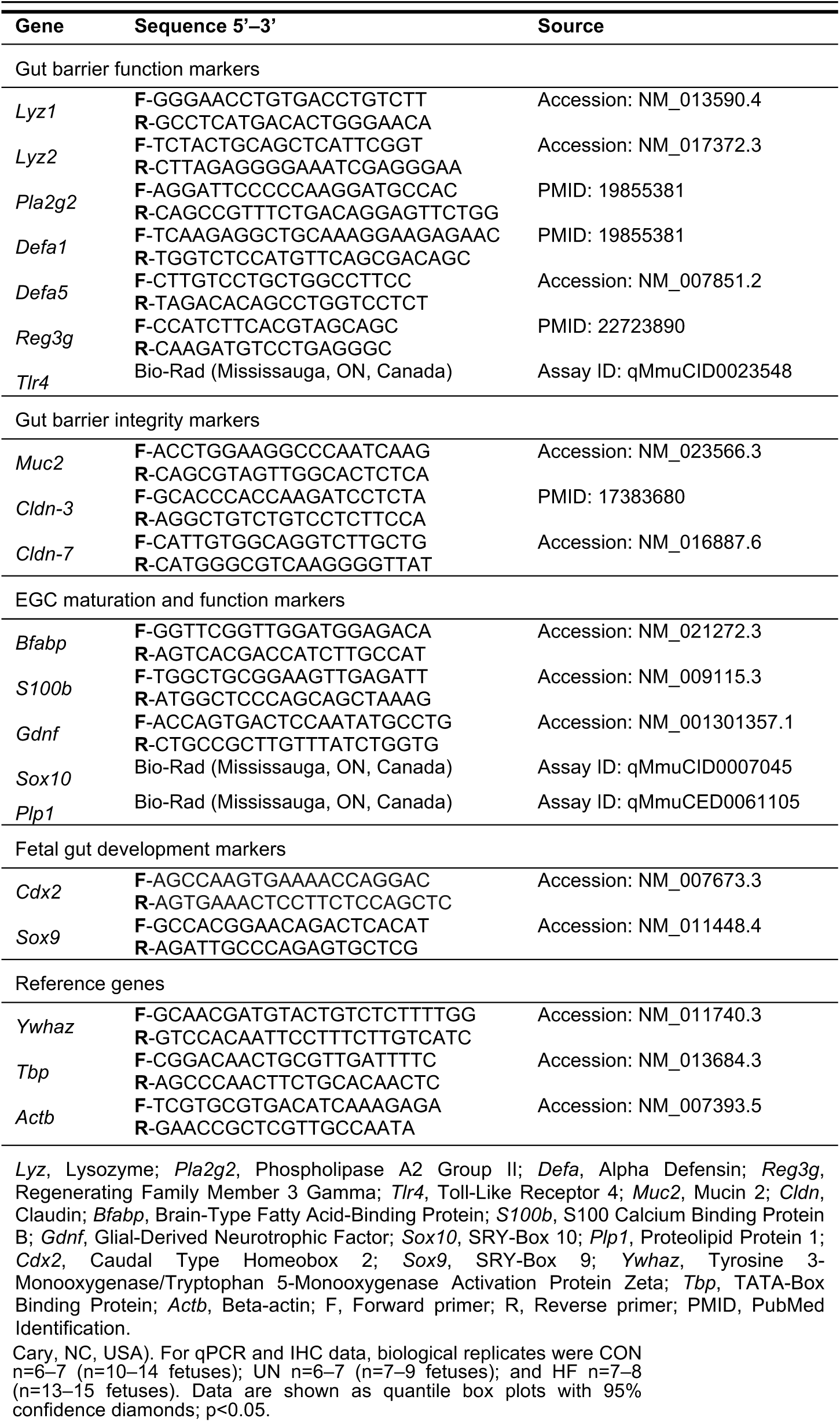
Primer sequences for quantitative PCR.

### Expression and localization of lysozyme protein and quantification of goblet cells in maternal SI

Immunohistochemistry (IHC) was used to localise and semi-quantify immunoreactive (ir) staining of lysozyme (Lyz) and to quantify goblet cell number. Five mm sections of maternal SI were cut from paraffin-embedded blocks and mounted onto glass slides. For lysozyme staining, sections were deparaffinised and rehydrated in descending alcohol series and quenched with 3% hydrogen peroxide in 90% ethanol for 20min at room temperature. Sections underwent antigen retrieval by sodium citrate/citric acid solution and microwaved for 10min, then blocked in serum-free protein blocking solution (Agilent Dako, Santa Clara, CA, USA) for 1hr at room temperature, and incubated overnight at 4°C with rabbit anti-lysozyme antibody (1:200 dilution; product #PA1-29680, Thermo Fisher Scientific, Waltham, MA, USA). Negative control sections were probed with normal rabbit IgG (0.4ug/uL, 1:200; #sc-2027, Santa Cruz Biotechnology, Dallas, TX, USA). Sections were then incubated with goat biotinylated anti-rabbit secondary antibody (1:200; #BA-1000, Vector Labs, Burlingame, CA, USA) for 1hr at room temperature, followed by 1hr incubation with streptavidin-horseradish peroxidase (1:2000 in 1X PBS; Invitrogen, Carlsbad, CA, USA). Antibody-antigen interactions were visualised using DAB for 40sec (Vectastain DAB ABC kit, Vector Labs). Sections were counterstained with Gills I haematoxylin to provide greater contrast and to stain nuclei. Semi-quantitative analyses of ir-lysozyme staining intensity was performed using computerised image analysis (Image-Pro Plus 4.5, Media Cybernetics, Rockville, MD, USA; and Olympus BX61 microscope, Shinjuku, Tokyo, Japan). A superimage was composited from individual images (at 20x magnification) captured along the entire SI section from each dam. To semi-quantitatively measure staining intensity in each image (where higher levels of staining intensity may represent higher Lyz protein expression), an algorithm was developed (using Visiopharm NewCAST Analysis software, Hørsholm, Denmark) to detect intensity of DAB staining for ir-Lyz in four levels of intensity: low, moderate, high, and strong. Staining intensity for each of the four levels was summed across all images within a superimage for each animal. CON mothers had a mean ± SEM (range) of 10.3 ± 0.7 (7–13) images per superimage, UN had 10 ± 0.8 (7–13) images, and HF had 9 ± 0.6 (7–12) images. There was no statistical difference between diet groups in the number of images per superimage (p=0.37).

For goblet cell quantification, sections were stained with alcian blue, which stains acetic mucins and acid mucosubstances. Sections were deparaffinised and rehydrated in descending alcohol series then incubated in 3% glacial acetic acid, followed by alcian blue (Sigma-Aldrich, Oakville, ON, Canada) and 0.1% nuclear fast red counterstain (Electron Microscopy Sciences, Hatfield, PA, USA). Eight images were randomly captured at 20X for each section (Leica DMIL LED inverted microscope, Wetzlar, Germany; and QCapture Pro software, Surrey, BC, Canada). We first counted the number of alcian blue positive cells (goblet cells) in one villus and one crypt nearest to the villus in each of the 8 images and took the average across the images to determine the mean number of goblet cells in a villus and crypt for each animal. We also counted the number of alcian blue positive cells in two villi and two crypts nearest to these villi in each of four images (randomly selected from the eight images). The total number of epithelial cells in the same villi and crypts were also counted. The number of alcian blue positive cells and total epithelial cells across the two villi and two crypts in each image were summed. Then the number of alcian blue positive cells and total epithelial cells were averaged across the four images (separately for villi and crypts) to obtain the average number of goblet cells and epithelial cells and the percentage of goblet cells per two villi or crypts for each animal. A researcher blinded to the experimental groups performed the counting.

### Data analysis

Data were checked for normality using the Shapiro-Wilk test and equal variance using Levene’s test. Outliers were excluded from analyses. Non-parametric data were transformed by applying either logarithmic, square root, or cube root transformations. Differences between dietary groups for outcome measures were analyzed using: 1) ANOVA with Tukey’s *post hoc*, 2) Kruskal-Wallis/Wilcoxon test with Steel-Dwass *post hoc* for non-parametric data, or 3) Welch’s test with Games-Howell *post hoc* for normal data with unequal variance (p<0.05) using JMP 13 software (SAS Institute, Cary, NC, USA). For qPCR and IHC data, biological replicates were CON n=6–7 (n=10–14 fetuses); UN n=6–7 (n=7–9 fetuses); and HF n=7–8 (n=13–15 fetuses). Data are shown as quantile box plots with 95% confidence diamonds; p<0.05.

## Results

### Malnutrition was associated with reduced gut barrier function and integrity

In UN mothers, SI mRNA expression levels of AMP genes *Lyz2* (p=0.02 Figure 1B) and *Reg3g* (p=0.003, Figure 1C) were decreased compared to CON. HF mothers had reduced mRNA expression levels of *Reg3g* (p=0.003, Figure 1C) and *Muc2* (p=0.001, Figure 1G) in SI compared to CON. There was no effect of maternal diet on the expression of AMP genes *Lyz1, Defa1, Defa5*, and *Pla2g2* (Figures 1A, D-F). The average number of goblet cells, sites of mucus secretion, was higher in SI villi, but not crypts, of UN mothers compared CON (Figure 2). Proportion of goblet cells relative to total number of epithelial cells in both villi and crypts were not different between dietary groups (Figure 2).

**Figure 1.**
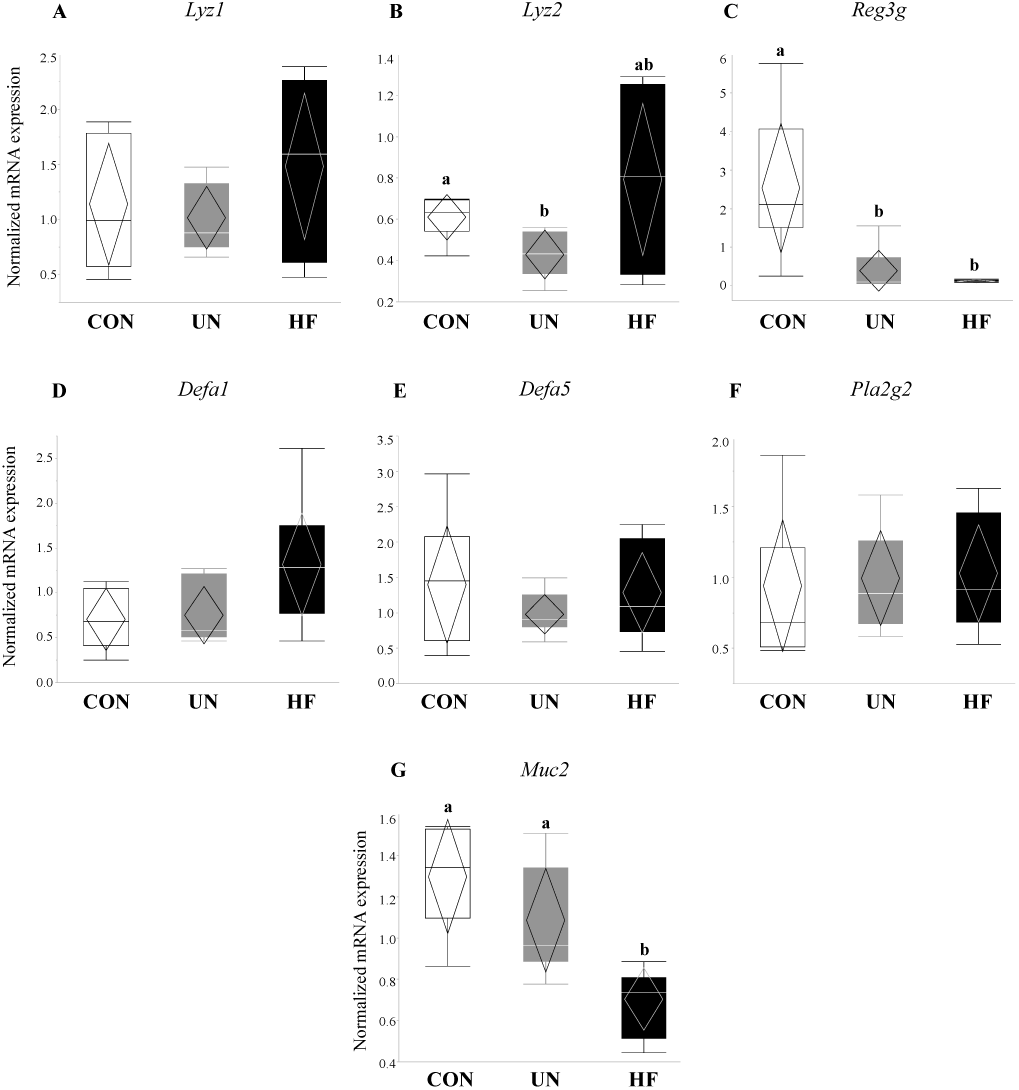
Maternal malnutrition was associated with altered gene expression of antimicrobial peptides & mucin. Maternal UN was associated with decreased mRNA expression of antimicrobial peptide genes *Lyz2* (p=0.02) and *Reg3g* (p=0.003) *vs.* CON, while HF diet was associated with decreased *Reg3g* (p=0.003) and mucin (*Muc2*; p=0.001) mRNA expression *vs.* CON (n=6–8/group). Groups with different letters are significantly different (p<0.05). UN, undernourished; HF, high-fat; CON, control.

**Figure 2.**
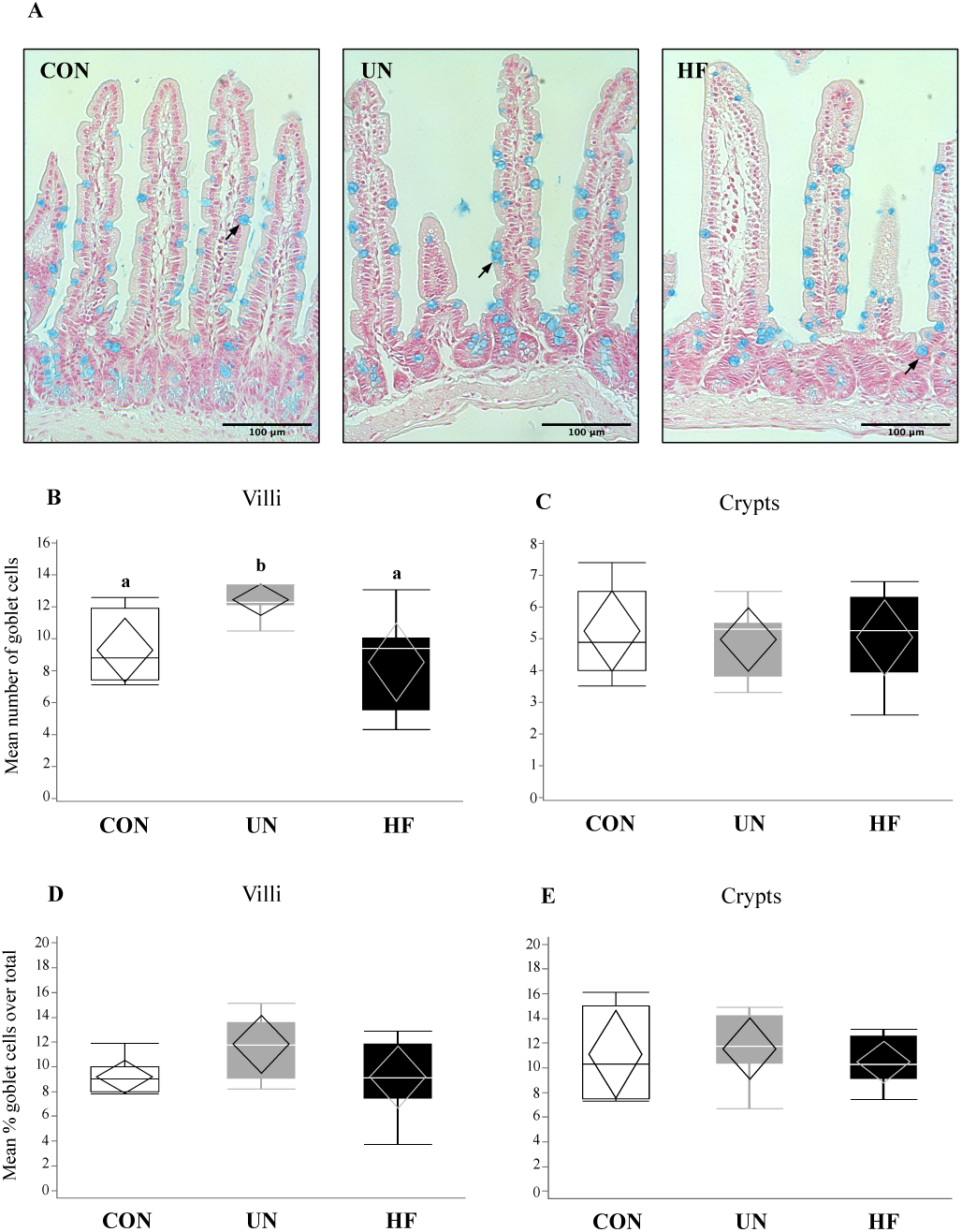
Maternal malnutrition may influence small intestinal goblet cell number. (A) Staining of goblet cells by alcian blue in small intestine (20X magnification). Arrows indicate goblet cells. Mean number of goblet cells across 8 villi (B) or crypts (C). Mean percentage of goblet cells (proportion of total number of epithelial cells) in villi (C) or crypts (D). There were a greater number of goblet cells in UN villi *vs.* CON (p=0.008; n=6– 8/group). Groups with different letters are significantly different (p<0.05). UN, undernourished; HF, high-fat; CON, control.

### Maternal HF diet may be associated with increased Lyz production

Lyz protein was localised to Paneth cells in the crypts of the maternal SI (Figure 3A). In the SI of HF dams, semi-quantitative analyses revealed a significant reduction in the levels of low-intensity Lyz staining (p=0.03, Figure 3B) compared to CON, which may suggest an overall greater production of this AMP in HF mothers. Further, an overall difference in moderate-intensity staining (p=0.04; Figure 3C) was detected, but there was no difference between groups with post hoc testing, and there were no differences between diet groups in high- or strong-intensity staining levels (Figures 3D-E).

**Figure 3.**
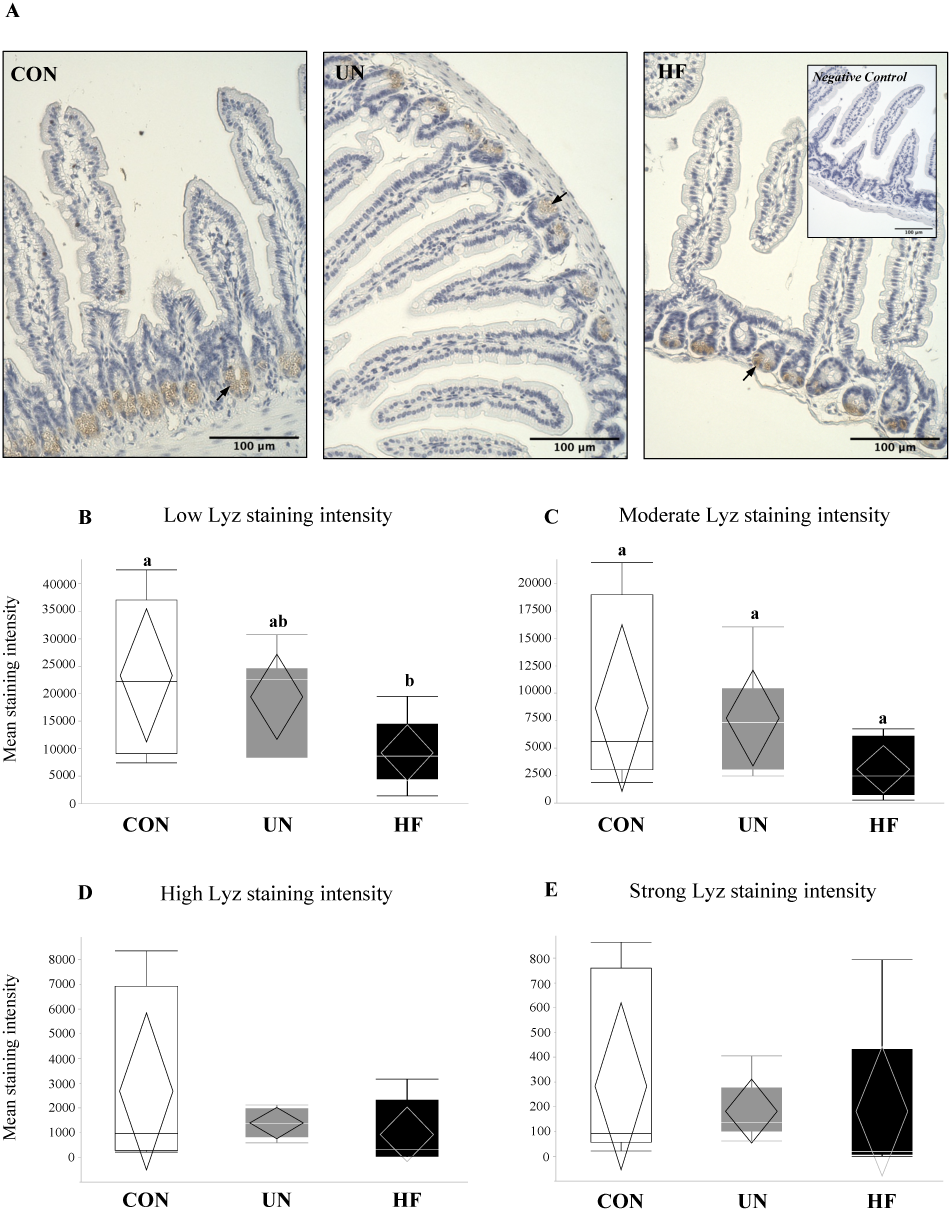
Maternal HF diet was associated with less low-intensity lysozyme staining in intestinal crypts. (A) Representative images of lysozyme protein immunoreactivity (ir) staining show localisation to the crypts of the maternal small intestine (SI) at d18.5, with negative control inset (40X magnification). Arrows indicate lysozyme proteins within Paneth cells. (B-E) Lyz staining was quantified into low, moderate, high, and strong intensities, representing increasing levels of protein expression (n=6– 8/group). Semi-quantitative analysis revealed less low-intensity Lyz staining in SI from HF mothers (p=0.03) *vs.* CON, and an overall difference in moderate-intensity Lyz staining (p=0.04), but no difference between groups with *post hoc* testing. Groups with different letters are significantly different (p<0.05). UN, undernourished; HF, high-fat; CON, control.

### Maternal UN was associated with delayed fetal gut development & reduced mucus production

UN, but not HF, fetuses showed increased mRNA expression of the gut transcription factor *Sox9* (p=0.02, Figure 4A) compared to CON, and an associated decrease in the mRNA expression levels of *Muc2* (p=0.002, Figure 4B) and *Cdx2* (p=0.003, Figure 4C) compared to CON.

**Figure 4.**
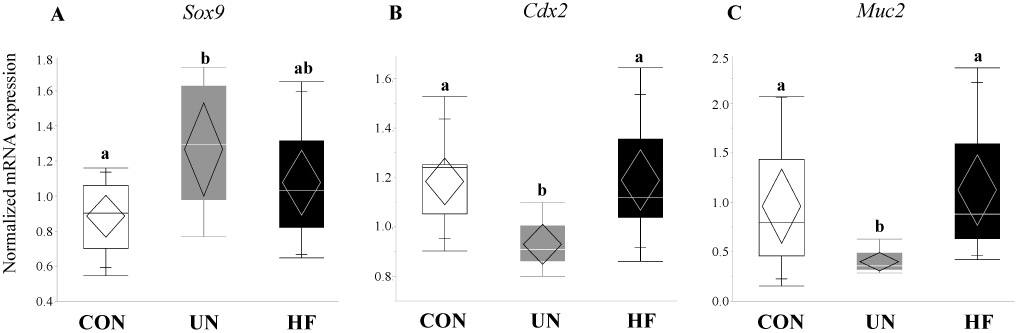
UN fetuses displayed activation of a gut transcription factor that represses gut barrier development and mucus production. Maternal UN was associated with increased fetal gut mRNA expression of Sox9 (p=0.02) vs. CON, and decreased Muc2 (p=0.002) and Cdx2 (p=0.003) vs. CON (n=8–15/group). Groups with different letters are significantly different (p<0.05). UN, undernourished; HF, high-fat; CON, control.

### Maternal HF diet was associated with increased fetal EGC development

Fetuses from HF-fed mothers had increased mRNA expression levels of EGC markers *Bfabp* (0.003, Figure 5A) and *S100b* (p<0.001, Figure 5B) compared to CON, and increased *Plp1* (p=0.04, Figure 5C) compared to UN. There were no differences in gut mRNA expression levels of the EGC marker *Sox10* and EGC neurotrophic factor *Gdnf* in fetuses exposed to different maternal diets (Figures 5D-E).

**Figure 5.**
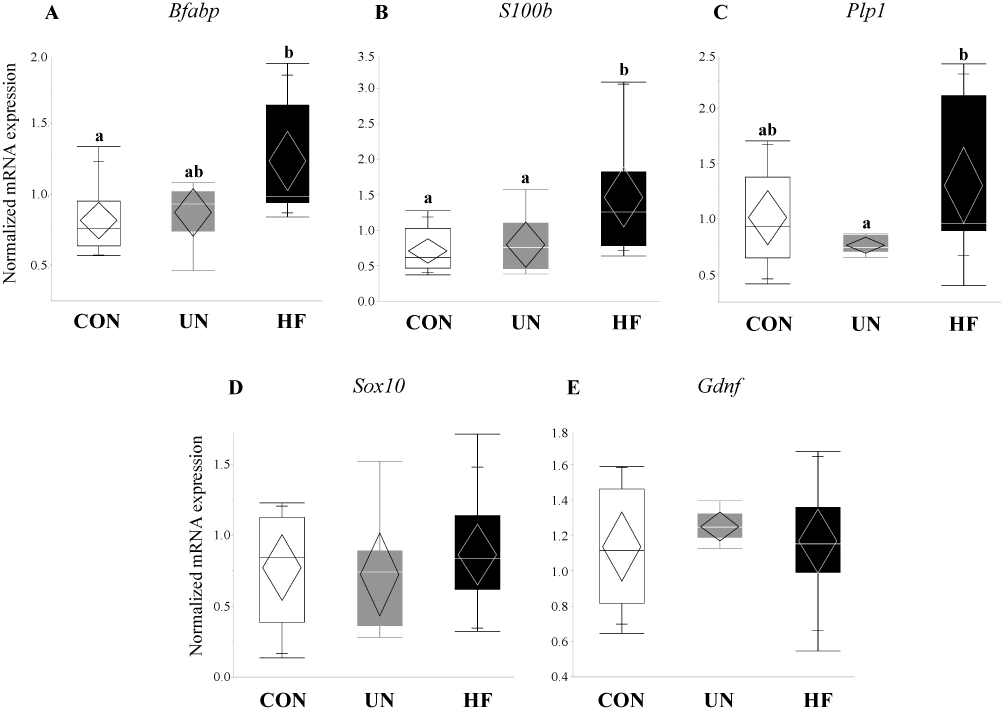
HF fetuses showed activation of enteric glial cell markers in the gut. Maternal HF diet was associated with increased fetal gut mRNA expression of EGC markers *S100b* (p<0.001) and *Bfabp* (p=0.003) *vs.* CON, and *Plp1* (p=0.04) *vs.* UN (n=7–15/group). Groups with different letters are significantly different (p<0.05). UN, undernourished; HF, high-fat; CON, control.

### Maternal HF diet was associated with increased fetal gut barrier function and integrity

HF, but not UN, fetuses had increased mRNA expression levels of Paneth AMPs *Lyz1* (p=0.007, Figure 6A), *Reg3g* (p=0.01, Figure 6C), and *Defa1* (p=0.001, Figure 6D), and lower gut mRNA expression levels of *Tlr4* (p=0.02, Figure 6E) compared to CON. No between-group differences were detected in *Lyz2* fetal gut mRNA expression levels (Figure 6B). Maternal HF diet was also associated with an increase in fetal gut mRNA expression of TJ genes *Cldn-3* (p=0.008, Figure 7A) and *Cldn-7* (p<0.001, Figure 7B) compared to CON, while UN fetuses only displayed increased mRNA expression of *Cldn-7* compared to CON (p<0.001, Figure 7B).

**Figure 6.**
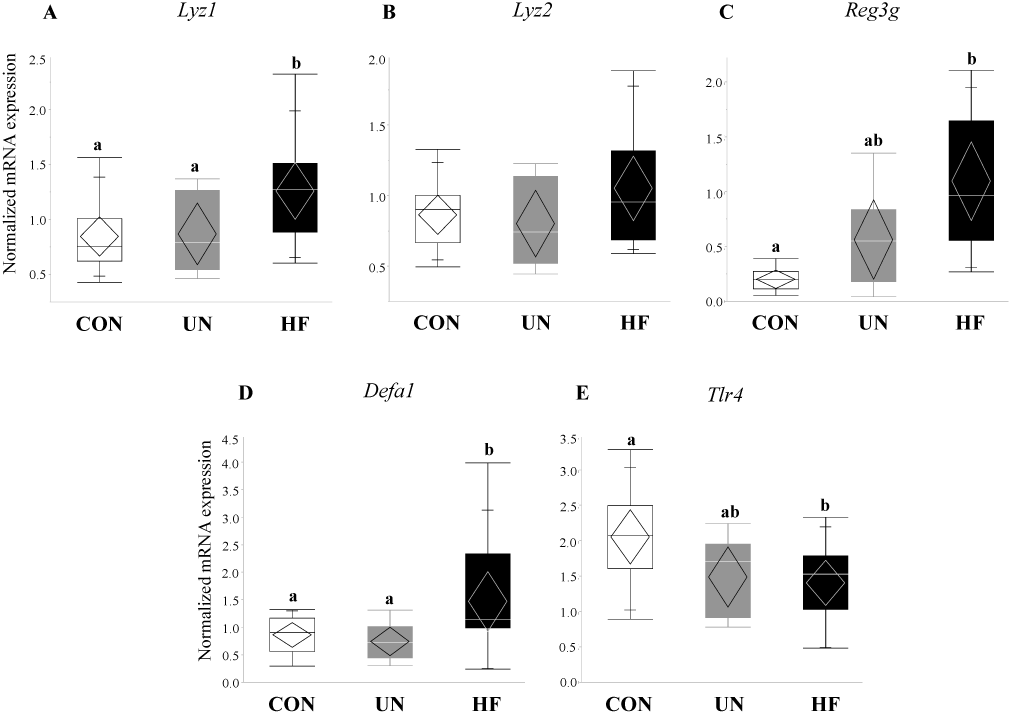
HF fetuses showed an upregulation of gut barrier function genes and downregulation of a microbe-sensing receptor. Maternal HF diet was associated with increased mRNA expression of antimicrobial peptide (AMP) genes *Lyz1* (p=0.007), *Reg3g* (p=0.01), and *Defa1* (p=0.001) in the fetal gut *vs.* CON, though mRNA expression of the purported AMP-activating receptor *Tlr4* was decreased in these fetuses (p=0.02) *vs.* CON (n=9–15/group). Groups with different letters are significantly different (p<0.05). UN, undernourished; HF, high-fat; CON, control.

**Figure 7.**
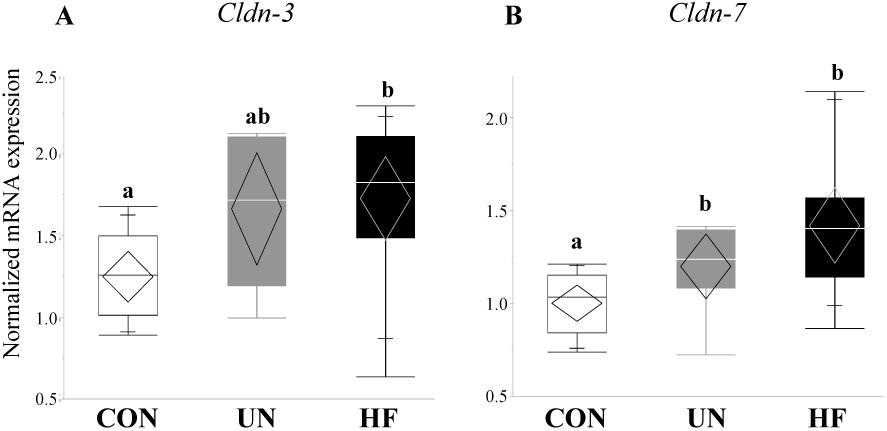
Maternal malnutrition altered fetal gut tight junction gene expression. Maternal UN was associated with increased mRNA expression of *Cldn-7* (p<0.001) in fetal gut *vs.* CON, while fetuses from HF mothers increased *Cldn-3* (p=0.008) and *Cldn-7* (p<0.001) mRNA expression *vs.* CON (n=9–15/group). Groups with different letters are significantly different (p<0.05). UN, undernourished; HF, high-fat; CON, control.

### Maternal malnutrition altered fetal gut mRNA expression in males more than in females

Data on fetal outcomes were also stratified by sex (Figures 8–11). We found that maternal HF diet was associated with increased mRNA expression of genes involved in EGC maturation (*Bfabp, Plp1*, and *S100b*), gut barrier function (*Lyz1* and *Lyz2*), and gut barrier integrity (*Cldn-3* and *Cldn-7*) in the male fetal gut (p<0.05, Figures 9–11) compared to CON. Maternal HF diet was also associated with reduced male fetal gut mRNA expression of the microbe-sensing toll-like receptor *Tlr4* (p=0.03, Figure 10E) compared to CON. Male UN fetuses showed a reduction in fetal gut mRNA expression of gut differentiation and maturation markers *Cdx2* (p=0.02, Figure 8B) and *Muc2* (p<0.001, Figure 8C), and an upregulation of EGC marker *Bfabp* (p<0.001, Figure 9A) and TJ gene *Cldn-7* (p=0.004, Figure 11B).

**Figure 8.**
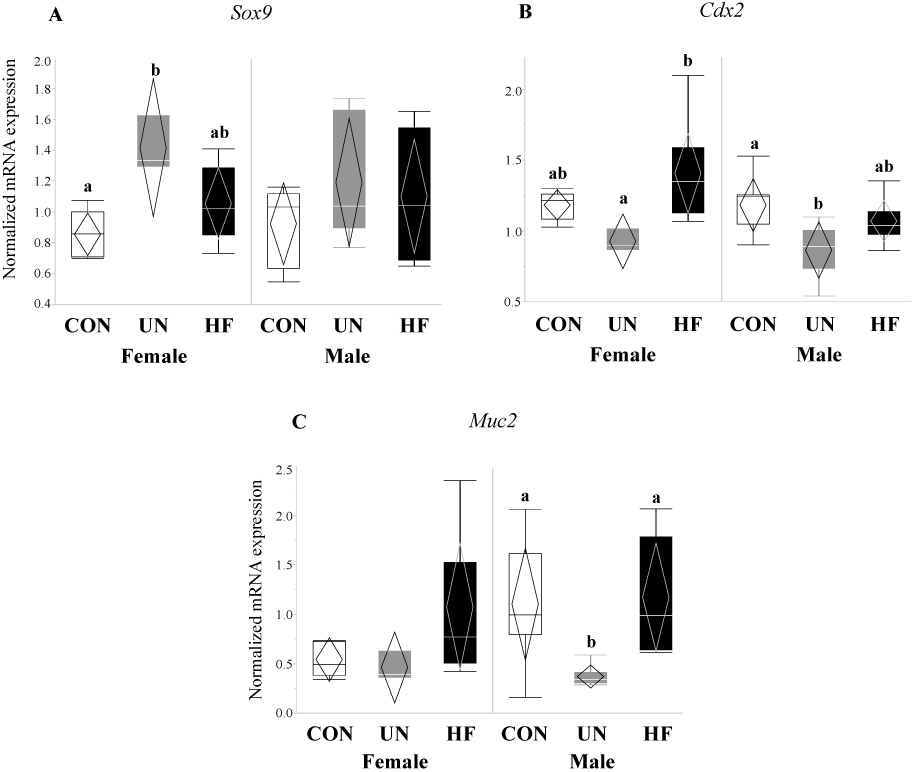
Maternal UN affected fetal gut barrier maturity in both sexes and mucus layer maturity in male fetuses only. Maternal UN was associated with increased gut transcription factor *Sox9* (p=0.004) *vs. CON* and decreased *Cdx2* (p=0.02) *vs. HF* in female fetal guts, and reduced *Cdx2* (p=0.02) and *Muc2* (p<0.001) *vs.* CON in male fetal guts (n=3– 8/group). Groups with different letters are significantly different (p<0.05). UN, undernourished; HF, high-fat; CON, control.

**Figure 9.**
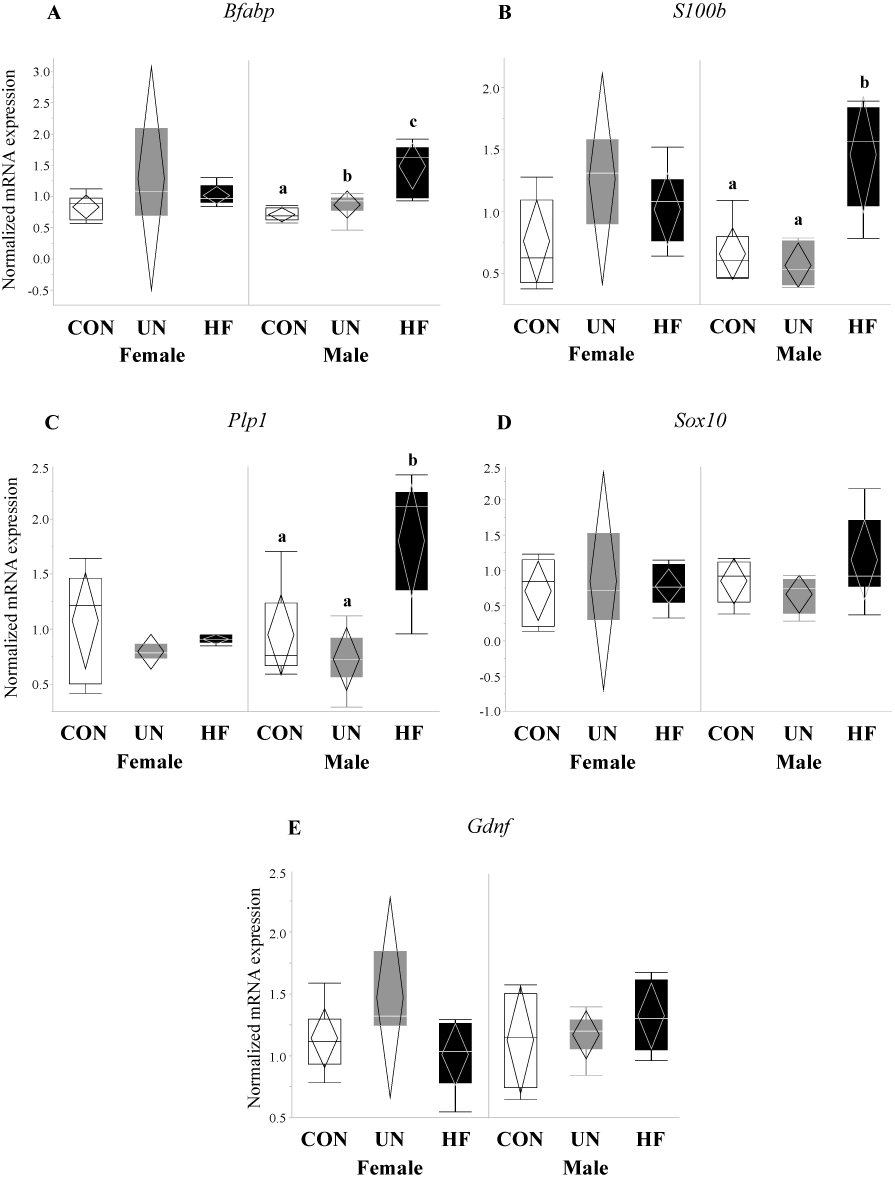
Maternal HF diet was associated with increased mRNA expression of enteric glial cell markers in male fetuses. In male fetal guts, maternal HF diet was associated with increased *Bfabp* (p<0.001), *S100b* (p<0.001), and *Plp1* (p=0.001), while maternal UN was associated with increased *Bfabp* (p<0.001). Maternal malnutrition (UN & HF) did not affect enteric glial cell development in female fetal guts (n=3–8/group). Groups with different letters are significantly different (p<0.05). UN, undernourished; HF, high-fat; CON, control.

**Figure 10.**
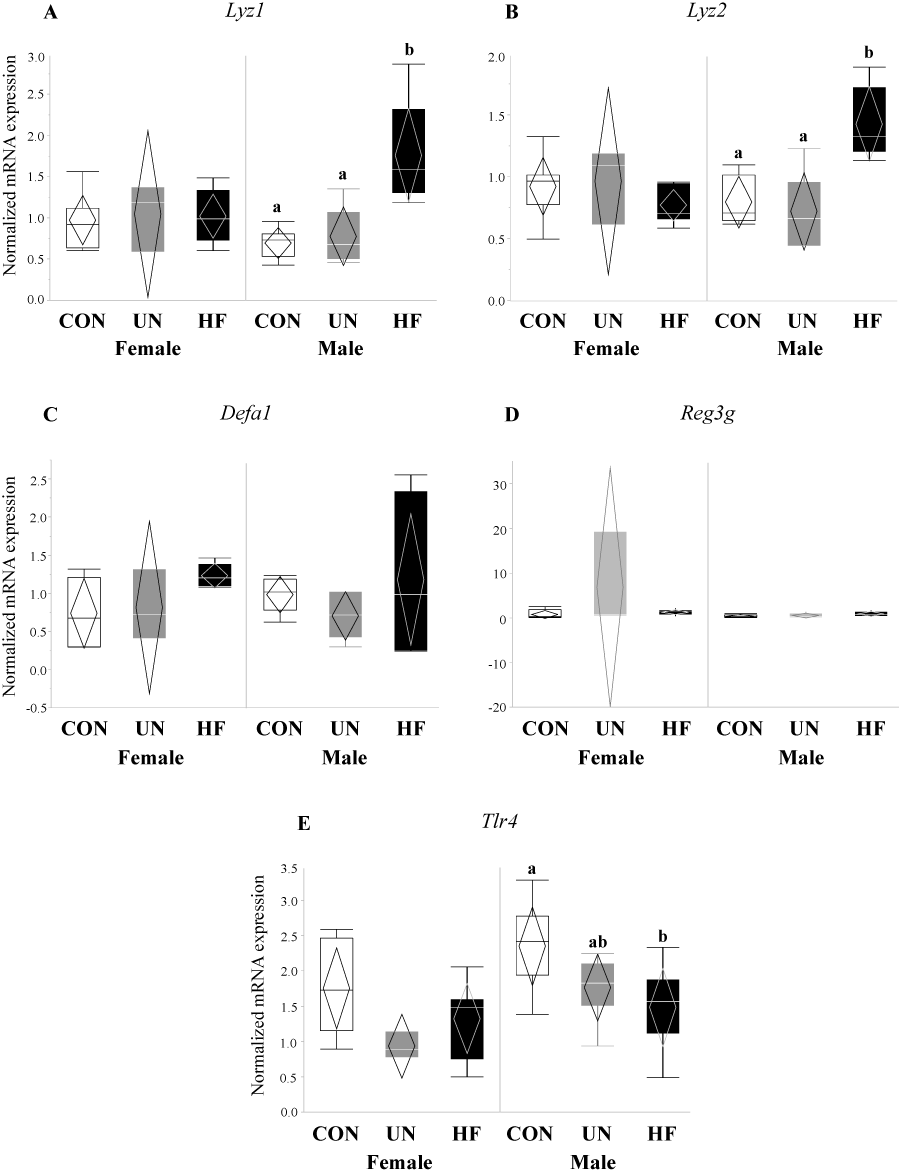
Maternal HF diet was associated with increased expression of antimicrobial peptides and decreased expression of microbe-sensing receptor in male fetuses. In male fetal guts, maternal HF diet was associated with increased *Lyz1* (p<0.001) and *Lyz2* (p<0.001), and decreased *Tlr4* (p=0.03) mRNA expression levels. Maternal malnutrition (UN & HF) did not affect *Defa1* or *Reg3g* expression levels in male fetuses or antimicrobial peptide levels in female fetal guts (n=3–8/group). Groups with different letters are significantly different (p<0.05). UN, undernourished; HF, high-fat; CON, control.

**Figure 11.**
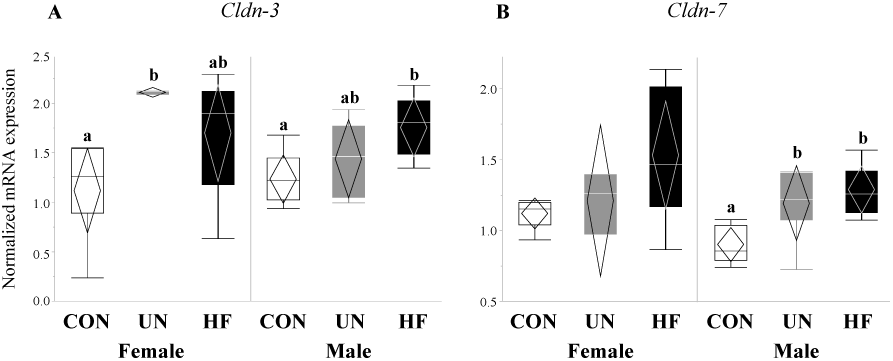
Maternal malnutrition was associated with an increase in gut tight junction gene expression in both fetal sexes. In male fetal guts, maternal HF diet was associated with increased *Cldn-3* (p=0.02) and *Cldn-7* (p=0.004), while maternal UN was associated with increased *Cldn-7* (p=0.004); in female fetal guts, maternal UN was associated with increased *Cldn-3* (p=0.03) mRNA expression (n=3–8/group). Groups with different letters are significantly different (p<0.05). UN, undernourished; HF, high-fat; CON, control.

In female fetal guts, maternal UN was associated with increased mRNA expression of gut differentiation transcription factor *Sox9* (p=0.004, Figure 8A) compared to CON, and decreased gut maturation marker *Cdx2* (p=0.02, Figure 8B), though only compared to HF. Maternal UN was also associated with increased TJ gene *Cldn-3* (p=0.03, Figure 11A) in female fetuses compared to CON.

## Discussion

Despite the growing body of evidence linking the gut and its resident microbes to health (2,6), few studies have investigated these relationships in the mother during pregnancy or the developing fetus. Additionally, although we know that key enteric cells that support the gut barrier and establish communication with the brain are laid down and functional in early life (54–56), we know less about how early life nutritional adversity or altered gut microbes may influence their development and function (57,58). Since malnutrition is an important insult to pregnancies and their outcomes (59), our study is the first to determine the effect of maternal malnutrition on maternal and fetal intestinal barrier integrity and function, and its implications for fetal development.

Extensive research has demonstrated that high-fat diets markedly alter the diversity and composition of gut microbes (40,60–64), resulting in long-term aberrations in gut barrier integrity (40,60–62,65) and function (40,60,66) and chronic inflammation (40,61,62). As a result, we hypothesised that mothers fed a HF diet would show pronounced changes in gut function. We found that maternal HF diet was associated with reduced mRNA expression of Paneth cell AMP *Reg3g*, which targets gram-positive bacteria (67), no change in mRNA expression of AMPs *Lyz1* and *Lyz2,* and less low-intensity Lyz staining, which may suggest an overall greater production of Lyz protein, which can target both gram-positive and gram-negative bacteria (68). These results are analogous to those where HF feeding for 8 weeks in non-pregnant mice was associated with a reduction in *Reg3g* SI mRNA expression and no expression changes in *Lyz1* (66). Maternal HF diet was also associated with lower mRNA expression levels of the goblet cell-produced *Muc2*, which when translated, becomes the main component of the mucus layer: the first line of defence against gut infections and inflammation (69). Although this was not associated with an increase in the number of goblet cells in HF SI, our mRNA results align with a study where male and female mice fed a HF diet from weeks six to 22 of life had a reduction in mRNA expression levels of *Muc2* and cryptidins (Paneth AMPs) (70). Since Reg3g is critical to Muc2 distribution and spatial segregation of the gut epithelium and bacteria (67,71,72), concurrent downregulation of these genes due to a HF diet may further alter the mucus layer and reduce gut barrier defences against microbes, resulting in bacterial contact with the epithelium, increased gut inflammation, gut tissue damage, and bacterial translocation (67).

Similar to high-fat diets, undernutrition is known to be a significant insult to gut barrier integrity and function, increasing gut permeability (73–75), gut and peripheral inflammation (73,74), and bacterial translocation (76), and altering gut barrier structure (77) and enteric cell function (78,79). Despite this, and to the best of our knowledge, our study is the first to investigate the effects of undernutrition on the maternal gut barrier during pregnancy. We hypothesised that undernutrition would lead to adverse changes in the maternal gut environment. We found that moderate maternal UN was associated with lower mRNA expression levels of Paneth cell AMPs (*Lyz2* and *Reg3g*), suggesting reduced gut barrier function and ability to maintain gut-microbe homeostasis. This is consistent with findings from a study where 48 hours of starvation in non-pregnant mice led to a reduction in SI mRNA and protein expression of Reg3g and Lyz, and a hyper-permeable barrier (41). Since AMPs limit bacterial contact with the gut barrier (80), a reduction in their expression has been associated with increased bacterial adherence to the barrier (81) and bacterial translocation (1,41,82). Moreover, AMPs have been shown to block the release of IL-1β from activated immune cells (83), a pro-inflammatory cytokine which exacerbates gut barrier permeability by creating gaps between TJ proteins (83–85). These data suggest that undernutrition may compromise the host’s ability to mount an appropriate immune response through AMP pathways, resulting in increased susceptibility to infections and a leaky gut. In fact, malnutrition is known to impair gut immune functioning, causing increased susceptibility to environmental insults due to altered cytokine production (86). This may be of particular concern in pregnancies where undernutrition/underweight and infection often coexist, such as in populations with low socioeconomic status (30,75). Yet, even prior to bacteria reaching gut barrier, a mucosal layer produced by goblet cells provides protection. We found that UN mothers had an increased number of villus-residing goblet cells, which may suggest an attempt to strengthen the gut barrier to offset the consequences of reduced AMP expression, such as the heightened propensity for leaky gut.

Though early life development is key to setting healthy trajectories throughout life (87), little is known about how perinatal events shape fetal gut development. Therefore, we were interested in the effect of maternal malnutrition on fetal gut and ENS development and function. We focused on the consequence of this nutritional adversity on genes involved in gut maturation and differentiation, EGC development, Paneth AMP production, and TJ formation. Consistent with our hypothesis, fetuses from HF mothers seemed to be most affected by an adverse nutritional exposure, as evidenced by significantly altered expression levels of nine of the 15 genes tested. HF diet was associated with changes in fetal EGC development, gut AMP production, and TJ expression. In fetuses from HF-fed dams, we found increased mRNA expression of specific and widely-expressed (16,22,88) markers of EGCs *S100b, Bfabp* and *Plp1*, the latter of which to our knowledge has not been previously examined in the mouse fetal gut. This increase in EGC markers may be in response to the higher inflammatory environment in HF pregnancies that we have previously described (49), since EGCs are important for regulating inflammatory pathways (89) and gut barrier integrity (19,89). During intestinal inflammation in adult animals, EGCs respond to pro-inflammatory signals and reverse inflammation-induced ENS damage (90) by driving enteric neuron death through nitric oxide production (49,51,52) and triggering ENS neurogenesis (91). Previously, EGC proliferation was shown to occur in infant rats whose mothers were fed HF diets perinatally, though mRNA and protein expression of local pro-inflammatory cytokines were unchanged at all postnatal timepoints examined (2, 4, 6, and 12 weeks) (43). Together with our results, this may suggest that maternal HF diet exposure *in utero* may reprogramme mechanisms in the fetus that establish gut-brain connections and communication and increase EGC development to reduce gut inflammation by the time of birth, thereby negating some of the fetal ENS damage incurred during development. Future experiments should examine whether these changes in the fetal gut are associated with altered brain development and function.

Additionally, fetuses from HF mothers showed an increase in gut mRNA expression levels of AMP genes *Lyz1* (but not *Lyz2*), *Defa1,* and *Reg3g*. This is in contrast to studies in adult mice (35,40,60,70) that have demonstrated that HF diets decrease gut levels of AMPs, though one study (37) found differing results between the mRNA and protein levels. Nevertheless, our study is the first to our knowledge to uncover how maternal malnutrition alters expression levels of AMPs in near-term fetuses and is the first to assess *Reg3g* mRNA expression in fetal tissues. Due to the lack of other developed immune mechanisms, fetuses may be increasing the expression of AMP genes to regulate and protect themselves from the pro-inflammatory fetal environment observed in HF pregnancies (92–95). In parallel to the upregulation of AMPs, we observed reduced expression of *Tlr4* mRNA in HF fetal gut, despite that, at least in adult models, *Tlr4* is known to initiate AMP production (96,97). Nonetheless, depending on tissue type, the same TLR can downregulate, upregulate, or not affect AMP production (98), adding to the complexity of the TLR-AMP relationship which is further compounded by the lack of work on TLR function in fetal gut tissue.

In mammals, 24 members of the claudin family exist, each with unique charge-selectivity preferences that dictate barrier permeability and tight junction structure and function (99). Of the claudin family, *Cldn-3* and *Cldn-7* are the amongst the most highly expressed in all sections of the mouse GI tract (100). *Cldn-3* has previously been shown to be affected by HF diets (65) and commensal bacterial colonization (101), while *Cldn-7* has been found to have functions outside of barrier integrity, such as maintenance of intestinal homeostasis (102), and both its mRNA and protein are highly expressed in the small and large intestine (102). Still, expression levels of neither *Cldn-*3 nor *Cldn-*7 have been recorded in mouse fetal guts. In our study, fetuses from HF mothers showed an increase in mRNA expression of both TJs *Cldn-3* and *Cldn-7*, however these data run counter to the effects of HF diets on TJ expression in adult mice and rats (61,62,65). Research on the ontogeny and development of tight junction proteins in the fetal mouse gut is scarce, though in human fetuses, tight junction proteins appear at 8–10 weeks of gestation and begin to assemble junctional complexes (continuous, belt-like structures around cells) at 10–12 weeks (103). Nevertheless, most of the development of the tight junction proteins and gut integrity occurs in postnatal life (99), indicating that the prenatal fetal gut may be particularly sensitive and vulnerable to the effects of adverse *in utero* exposures due to a permeable gut barrier. As HF fetuses showed increased expression of TJ proteins, which is consistent with their increased expression of EGC markers, it may be that HF fetuses are compensating for a highly inflammatory maternal environment by tightening the gut barrier.

Similar to HF fetuses, we hypothesised that fetal growth restriction would lead to aberrant gut barrier development in fetuses from UN mothers. Accordingly, we found that UN fetuses had increased expression of the gut transcription factor *Sox9*, which is known to repress the expression of *Muc2* and *Cdx2* through activation of the Wnt-β-Catenin-TCF4 pathway (104), and reduced mRNA expression of mucus (*Muc2*) and gut differentiation (*Cdx2*) genes. These data are indicative of immature gut barrier development, and consistent with the reduced weight of these fetuses compared to CON and HF fetuses. Importantly, an immature gut may be associated with increased susceptibility to inflammatory and infectious insults, due to reduced gut barrier integrity and function, gut dysbiosis (105–107), and irreversible (108) aberrant nutrient absorption (108–111). This has long-term implications for growth-restricted offspring, including those born too soon, such as preterm infants, who are at greater risk for necrotizing enterocolitis (112–114) and nutrient malabsorption (108,110,111) due to poor gut development and function. Despite the changes in gut maturation and mucus production, fetuses from UN mothers showed an increase in gut *Cldn-7*, but not *Cldn-3*, mRNA expression, which might suggest an attempt to increase barrier integrity to compensate for the gut barrier immaturity. Lastly, our sex-stratified data suggest that the majority of observed group differences were driven by male fetuses, which is consistent with numerous reports which demonstrate that male fetuses are more susceptible to perinatal insults, especially due to maternal malnutrition (115–120).

One limitation of our study is our focus on gene expression, especially in fetal samples where sample volume is extremely limited. Findings from qPCR data can direct future experiments that examine changes at the protein level. Another limitation is the low n-number in the female fetal UN group; thus, sex-stratified changes in mRNA expression should be interpreted with caution. Lastly, although our study is cross-sectional, we focused on an important developmental time (d18.5) that can serve as an indicator of embryonic/fetal experiences and provide information that could explain neonatal development and adaptations. Future studies should examine when changes in Paneth cell and EGC development and function are initiated in the pregnant mother and developing offspring, and how long they persist, which could point to critical windows for intervention to correct adverse health and developmental trajectories set by malnutrition.

## Conclusions

Our study is the first to examine the effect of both over- and undernutrition in parallel, during gestation in mice, and reveal the impact of maternal malnutrition on fetal gut Paneth cell function and EGC development — cells vital for gut barrier function and gut-brain axis connection. Our results indicate that malnutrition before and during gestation has adverse consequences for fetal gut development, maternal and fetal gut function, and potentially long-term programming of gut and brain function and gut immunity. If our findings are applicable to humans, this work may help inform research on dietary interventions that aim to prevent or mitigate the effects of being exposed to suboptimal nutrition in pregnancy (for mothers) and in early life (for offspring). Future work should place special focus on vulnerable populations wherein malnutrition and infection are more likely to coexist during pregnancy, exacerbating the negative repercussions of both noxious states on maternal and offspring wellbeing.

## Acknowledgments

We deeply thank Professor Stephen Lye for resource contributions towards the animal model; Tina Tu-Thu Ngoc Nguyen for the Lyz image capture and data collation; Ryszard Bielecki for the Lyz algorithm set up; and Enrrico Bloise for the goblet cell quantification. This research was funded by the Faculty of Science, Carleton University and the Canadian Institutes of Health Research (animal model MOP-81238)

## Author Contributions and Notes

Conceptualization, methodology, KLC; data curation, formal analysis, KLC and SAS; investigation, KLC, SAS, TTNN, EB; writing—original draft preparation, SAS; writing—review and editing, SAS and KLC; supervision, project administration, resources, KLC. The authors declare no conflict of interest.

---

**Figure S1.**
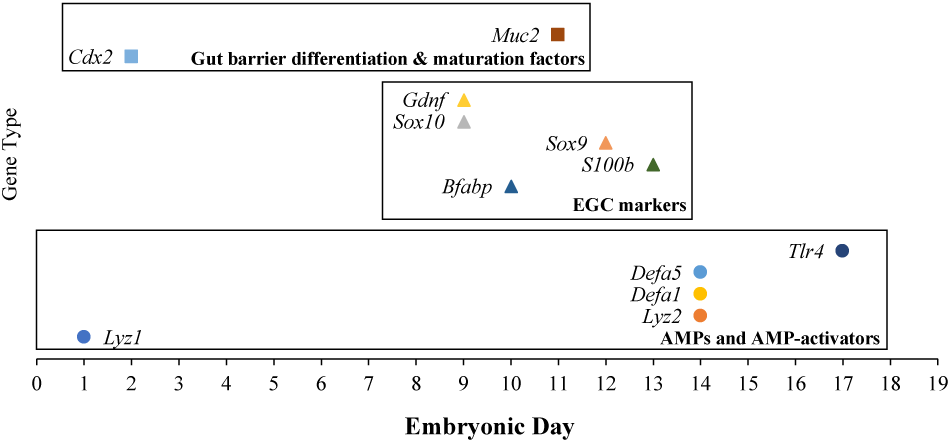
Earliest recorded expression of gut integrity, function, and development genes in the mouse fetal gut. Embryonic day of first established fetal gut expression of Paneth AMPs (*Lyz1, Lyz2, Defa1,* and *Defa5*), AMP-activator receptor (*Tlr4*), EGC markers (*Sox9, Sox10, S100b, Bfabp,* and *Gdnf*), gut barrier differentiation (*Cdx2*), and mucus production (*Muc2*) genes. Genes are grouped by function. Fetal gut expression levels of *Pla2g2, Reg3g, Cldn-3, Cldn-7,* and *Plp1* could not be found in the literature. Data from (13,22–29).

